# Learning Residual-based Biomarkers of Cognitive Health via Self-Supervised Learning on EEG State Transitions

**DOI:** 10.1101/2025.09.21.677598

**Authors:** Thomas Tveitstøl, Mats Tveter, Christoffer Hatlestad-Hall, Hugo L Hammer, Ira R J Hebold Haraldsen

## Abstract

Deep learning (DL) models have achieved impressive performance in EEG-based prediction tasks, but they often lack interpretability, limiting their clinical utility. In this study, we introduce a novel self-supervised learning (SSL) framework inspired by neurophysiological reactivity. Our approach models healthy EEG transitions between ocular states by predicting an EEG-derived feature under eyes-open conditions, using features from eyes-closed recordings. The residual between observed and predicted values quantifies deviations from normative brain dynamics and serves as a candidate biomarker. To improve clinical relevance, we propose two optimisation strategies that promote residuals predictive of pathology. We evaluated the framework using healthy cohorts (LEMON, Dortmund Vital) for self-supervised training, and the AI-Mind cohort for downstream prediction of plasma p-tau217 levels, a proxy for cognitive pathology. Despite extensive hyperparameter optimisation, predictive performance remained poor across all methods, including baseline models, suggesting limitations in the downstream proxy or input signal. Nonetheless, our approach provides a transparent methodology for transition-based EEG biomarker discovery grounded in self-supervised learning.

## 1. Introduction

Artificial intelligence (AI) is expected to play an increasingly important role in healthcare and medicine [1]. The AI-Mind project aims to develop predictive tools for identifying individuals with mild cognitive impairment (MCI) who are at risk of progressing to dementia [2]. Electroencephalography (EEG) provides valuable information for such models, as it captures electrical brain activity at the synaptic level with high temporal resolution [3]. Furthermore, deep learning (DL), a subfield of AI that processes data through multiple non-linear layers, is of high interest due to its strong ability to learn complex patterns in a datadriven manner, also beyond clinically recognised features [4].

DL has demonstrated strong performance in pattern recognition, with significant advancements in natural language processing, computer vision, and speech recognition [5, 6, 7, 8]. Moreover, applying DL to EEG data has shown promising results for predictive modelling [9, 10, 11, 12]. However, from a neurophysiological perspective, factors beyond empirical risk minimisation are important. Although DL models can extract complex patterns from EEG, their outputs are often not biologically interpretable. Developing DL methods that yield interpretable biomarkers is therefore an interesting research direction.

To address this, we propose a self-supervised learning (SSL) framework that models normative transitions between EEG ocular states - an approach inspired by *reactivity*, a well-established neurophysiological phenomenon related to arousal fluctuation, intrinsic attention allocation, and visual stimuli exposure [13, 14, 15, 16]. Specifically, our framework learns normative relationships between resting-state EEG during eyes-closed and eyes-open conditions. Distinct from common SSL-based approaches, we model expectation values of neural activity rather than learning latent feature representations. Furthermore, our approach addresses additional aspects beyond empirical risk minimisation, as we estimate expected neural responses under healthy conditions and compute residuals. The utilisation of these residuals, which represent deviations from healthy expectation values, results in a more transparent and neurophysiologically grounded output. Therefore, our framework builds on the neurophysiological concept of reactivity, which refers to characteristic transitions in brain activity that occur in response to stimuli, such as when a subject opens or closes their eyes [17].

In SSL, the DL model is trained on a *pretext task*, using *pseudo-targets* which are generated from the data itself rather than human-collected targets [18, 19]. An example from computer vision is the jigsaw-puzzle, where the model learns to identify the relative positioning of shuffled image tiles [20]. Several SSL techniques have been explored for EEG analysis. These include signal transformation recognition, where the pretext task requires to identify the transformation applied to the signal [21, 22, 23, 24, 25]. Masked graph autoencoders have been investigated with channel masking [26, 27] and frequency masking [26]. Furthermore, contrastive learning has been used with various data augmentation techniques [28, 29, 24, 30, 31, 10]. Masked temporal and spatial recognition were employed in [32], where Gaussian noise was added to an event related potential (ERP) component or a brain region as part of the pretext tasks. In [33], the model was trained to determine whether the EEG windows had been rearranged, while in [34], the pretext task involved predicting the next EEG slice. While many SSL-based approaches aim to produce general-purpose feature representations, some studies have tailored their pretext tasks to specific downstream applications. For instance, tailoring the pretext task to the downstream task was made on sleep EEG [31], as the design of the pretext task required an understanding of the sleep cycle. In a particularly comprehensive evaluation, demonstrated the effectiveness of SSL in sleep staging in the few-labels regime. Furthermore, a review on deep representation learning for brain-computer interfaces (BCI) identified the main motivations for its use, including performance improvement, learning robustness to noise, learning invariances, learning in small-data regimes, integrating heterogeneous components, and uncovering underlying structures in EEG data [36].

While current SSL methodology has proven effective for its purposes, we argue that there are additional avenues to explore for their application to EEG. The key distinction of our framework compared to previous works, is that, rather than learning latent vector representations or general-purpose embedding vectors, we aim to model expectation values relevant to cognitive health, and use the residuals as biomarkers. The key contributions of this study are:

1. We propose a novel SSL-based framework that learns normative EEG response patterns across ocular states and uses the deviation (residual) from these learned expectations as a potential biomarker.
2. We introduce two optimisation strategies which explicitly promotes residuals that carry relevant information rather than strict empirical risk minimisation on the pretext task. We highlight that these strategies are not bounded by our application, but can readily be applied to any residual-based biomarker, such as the brain age gap.
3. We perform transparent and well-documented testing of our designed methods through comprehensive hyperparameter optimisation.

## 2. Methods

This section is structured as follows: First, we introduce a generalised SSL-based framework for learning cross-ocular state expectation values on EEG data. Second, we propose two methods for deriving maximally relevant residuals by (1) supervised hyperparameter optimisation (HPO) and (2) semi-supervised multi-task learning (MTL). Third, we describe the specific pretext tasks used in our experiments, and outline additional experimental details such as datasets, hyperparameters (HPs) and HP distributions (HPDs), and baselines. All code is publicly available at https://github.com/thomastveitstol/SSL-Features-EEG.

### 2.1 Learning normative cross-ocular EEG relationships via SSL

Our framework is conceptually simple: predict a chosen scalar feature of the EEG in one ocular state using selected EEG features from the other ocular state. This serves as the pretext task. Then, this residual is used for predictive modelling on a downstream task of interest. Figure 1 illustrates this framework.

**Figure 1.**
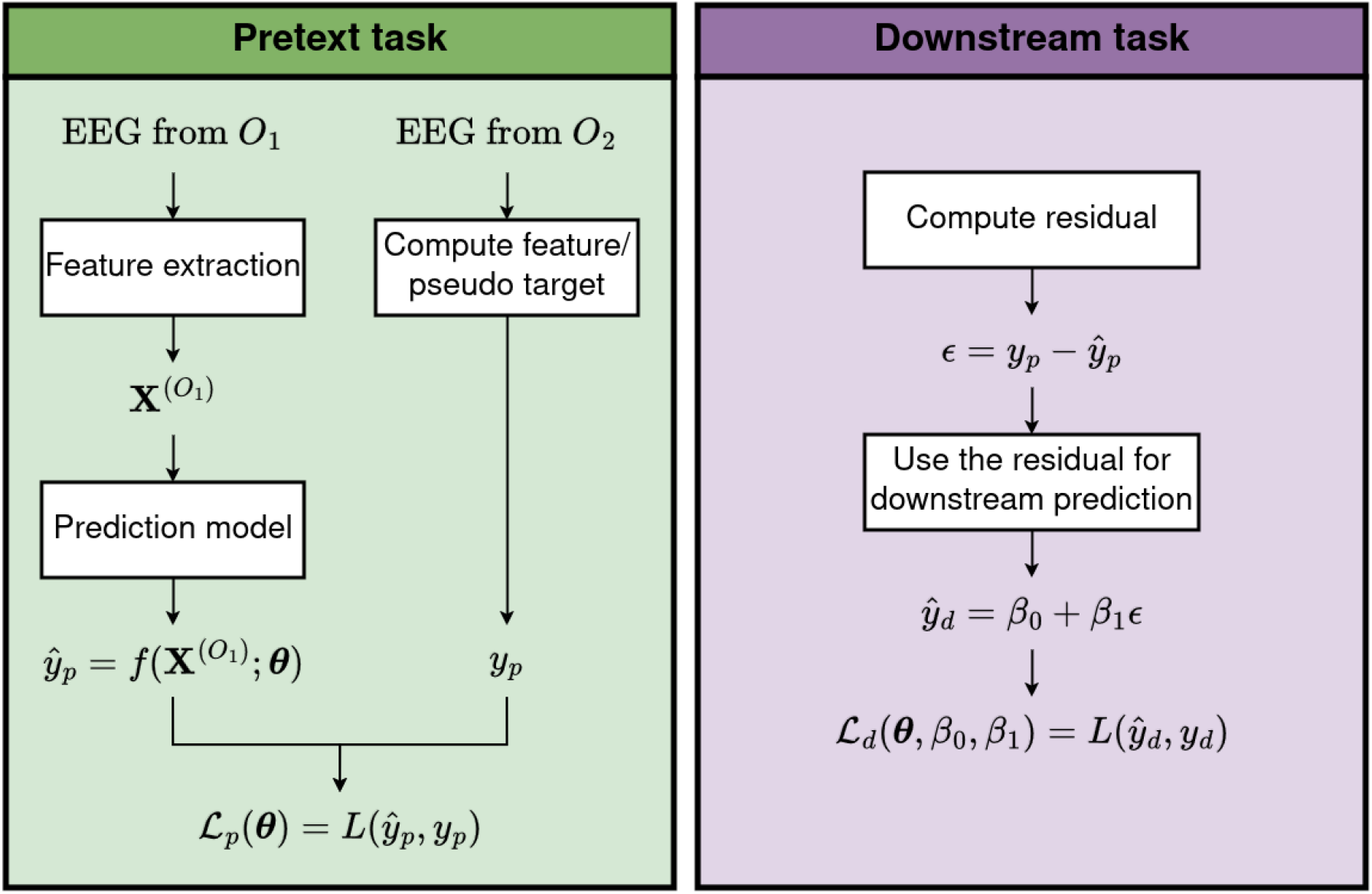
Overview of the proposed framework. In the pretext task, features 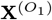 are extracted from the EEG in the ocular state *O*_1_, and used to predict a scalar feature/pseudo target, *y*_*p*_, calculated from the EEG in the ocular state *O*_2_. In the downstream task, the residual *ϵ*, which represents a deviation from the expectation value, is used for predictive modelling. While our figure illustrates a linear regression model, this approach can be extended to other machine learning models.

The first design choice in our framework is to select which ocular state should serve as input to the DL^2^ model, *O*_1_ ∈ {EC, EO }, and which ocular state the DL model will aim to predict a feature of, *O*_2_ ∈ {EC, EO} ^3^. To clarify, the EEG features from *O*_1_ are used to predict a scalar EEG feature derived from *O*_2_.

The EEG features, 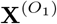, may focus on specific parts of the signals, such as a given frequency band through band-pass filtering, regions of interest through channel selection, the covariance matrix, or the EEG itself. In parallel, a scalar feature of choice is computed from the EEG during *O*_2_, denoted *y*_*p*_ ∈ ℝ. This scalar feature will serve as the pseudo-target of the pretext task. The core idea is that by designing and training a model to predict *y*_*p*_ from 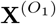, with 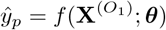, intuitive expectation values can be learned. Finally, the residual between observed and predicted value, *ϵ* = *y*_*p*_ − *ŷ*_*p*_, represents the deviation from expectation, which is hypothesised to reflect neurophysiologically meaningful deviations from normative brain responses and can be used as an interpretable biomarker for pathology prediction.

The isolated objective of the pretext task is to solve the following optimisation problem:

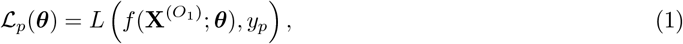

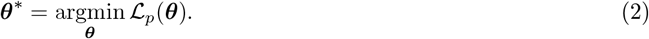

where ***θ*** is the model parameters.

The selection of the features 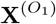 and pseudo target *y*_*p*_ determines the interpretation of the learned expectation value. For example, if the feature extraction involves computing the beta-filtered signal from frontal channels during the EO state, and the pseudo target is the correlation between the alpha-filtered channels Pz and C3 during the EC state, the expectation value represents: “Given EEG data from beta-filtered frontal channels during eyes open, what is the expected alpha correlation between Pz and C3 during eyes closed?”.

### 2.2 Optimising the predictive value of the residuals for pathological decoding

The main principle of our framework is to generate residuals that are both interpretable and useful for predictive modelling in a selected downstream task. This makes the modelling of *f* (·) and optimisation of ***θ*** in Eqs. 1 and 2 less straightforward, as in practice, the aim is not to minimise the residuals but to ensure that they carry relevant information. Since maximising performance on the pretext task does not necessarily increase the clinical utility of the residuals, we propose two methods for explicitly learning such meaningful residuals:

1. Supervised HPO, where the signal to the HPO algorithm is the predictive value of the residuals. This is quantified by how well a machine learning (ML) model can predict the downstream target using the residual. While we employed linear regression in our experiments and provide further details on its implementation in this section, this approach can be extended to other models.
2. Semi-supervised MTL where the first task is to predict the pseudo-target and the second task is to predict the downstream target from the residual.

The key difference between optimisation method 1 and 2 is that method 1 optimises the HPs to yield predictive residuals, while method 2 can also leverage gradient-based multi-task/multi-objective optimisation algorithms.

#### 2.2.1. Supervised HPO for maximising the predictive value of the residual

In supervised HPO, Eq. 2 is directly optimised for each trial during HPO, but model selection and the signal to the HPO algorithm is based on the predictive value of the residual. That is, we consider a predictive model *f* (·; **h, *θ***), where **h** ∈ ℋ denotes an HPC, and ℋ= *H*_1_ × · · · × *H*_*n*_ represents the HP configuration space. The HPO algorithm aims to find the optimal HPC **h**^∗^ by solving

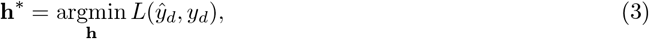

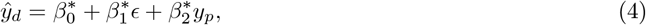

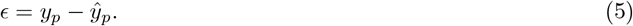

Here, 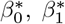, and 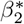 are the regression coefficients fitted by ordinary least squares after training on the pretext task (Eq. 2). The term containing the pseudo-target is included to account for any predictive signal in *y*_*p*_ itself. Without this adjustment, the HPO process might favour HPCs that produce constant predictions for *ŷ* _*p*_, causing the residual *ϵ* to trivially replicate the pseudo-target.

For HPO algorithms that iteratively refine HPCs based on prior performance, such as tree-structured parzen estimator (TPE) [37], supervised HPO can be implemented by passing linear regression performance scores such as *MSE, MAE*, or *R*^2^ to the HPO algorithm.

#### 2.2.2. Semi-supervised MTL for maximising the predictive value of the residual

A limitation of the optimisation method introduced in Sec. 2.2.1 is that it is restricted to only fitting the HPs to find residuals of high predictive value for the downstream task. An alternative method is to formulate our problem as a multi-task learning/multi-objective optimisation problem, where the first task is to solve the pretext task ℒ_*p*_ (Eq. 2), and the second task is to minimise the error on the downstream task ℒ_*d*_. Gradient-based algorithms from MTL and multi-objective optimisation may be applied to jointly optimise these two tasks simultaneously.

To impose our framework of making downstream predictions from learned residuals, the two objectives are formulated as

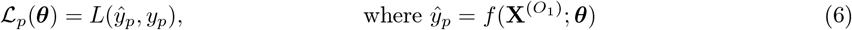

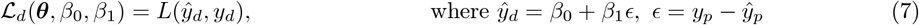

By jointly optimising these objectives, we are explicitly trying to learn the transition trends which are maximally relevant for pathology decoding.

We implemented five gradient based methods for MTL, which were (1) equal weighting of the loss functions, (2) GradNorm [38], (3) multiple-gradient descent algorithm (MGDA) [39, 40], (4) PCGrad [41], and (5) uncertainty weighting [42]. Below, these methods are briefly explained.

#### Equal weighting

The simplest multi-task learning strategy is to weigh the losses of the tasks equally. This was implemented by using the sum of the pretext and downstream loss, ℒ_*tot*_ = ℒ_*p*_ + ℒ_*d*_, and effectively treat the problem as a single-task optimisation problem.

##### GradNorm [38]

GradNorm applies gradient normalisation by dynamically altering gradient magnitudes. It was motivated by task imbalances and task dominance manifesting as imbalances in the magnitude of the backpropagated gradients. It uses adaptive weighted summation on the form *L*_*tot*_ = ∑_*i*_ *w*_*i*_*L*_*i*_, where *w*_*i*_ changes per training step, and the correct balance is obtained when the tasks are trained at similar rates [38]. As proposed in the original work, GradNorm was only applied on the final shared layer.

##### MGDA [39, 40]

MGDA is a gradient based and multi-objective optimisation based algorithm which aims to find a Pareto optimal solution. In MGDA, gradients are first applied on task specific parameters, followed by finding a direction of common descent, and applying this solution to the shared parameters. As the number of tasks in our study was equal to two, ℒ_*p*_ and ℒ_*d*_, we did not rely on the application of the Frank-Wolfe algorithm, and directly applied Algorithm 1 in [40] instead.

##### PCGrad [41]

PCGrad is an algorithm designed to mitigate conflicting gradients across tasks by preventing interfering components to be applied to the gradient-based parameter update. When two gradients are in conflict, a *gradient surgery* step is applied by projecting the gradients of each task onto the normal plane of the other.

##### Uncertainty weighting [42]

The uncertainty weighting approach developed in [42] weights the loss functions by using homoscedastic uncertainty. By formulating the problem as maximum likelihood estimation, it introduces a learnable noise parameter per task to adaptively learn the relative weighting of each loss based on task uncertainty.

### 2.3. Experiments

#### 2.3.1. Pretext tasks and downstream task

In this study, we considered pretext tasks aimed at learning the expected log band power during the eyes-open condition, given pre-processed EEG data from the eyes-closed condition. The frequency range used to compute band power was treated as a categorical HP, with the following options: *δ* (1–4 Hz), *θ* (4–7 Hz), *α* (7–12 Hz), *β* (12–30 Hz), and *γ* (30–45 Hz).

We used the downstream target as the log-transformed concentration of tau phosphorylated at threonine 217 (p-tau217), which was selected as a surrogate target variable due to its established association with cognitive decline and dementia development [43].

#### 2.3.2. Data

For training on the pretext tasks, we used resting-state EEG data from the LEMON dataset [44] and the Dortmund Vital study [45, 46]. For the downstream task, we used resting-state EEG data and p-tau217 levels from the AI-Mind study [2]. We used three epochs for each participant, with varying input length (see Sec. 2.3.5). Below, we provide a brief overview of these datasets; full documentation can be found in the original works. Note also that the choice of using LEMON, Dortmund Vital, or both, for training on the pretext task was not predetermined. Instead, dataset selection for the pretext task was treated as an HP to optimise.

##### LEMON

We used the pre-processed version of the LEMON dataset [44], which contains EEG data from *N* = 203 healthy participants. Data collection took place in Leipzig, Germany, using a 62-channel active ActiCAP electrode system. Since the eye electrode was not used in this study, we retained 61 scalp electrodes, following the extended international 10-20 system. Missing channels were interpolated using the “MNE” interpolation method in MNE-Python [47].

##### Dortmund Vital

The dataset from the Dortmund Vital study [45, 46, 48] contains EEG data from *N* = 608 participants. However, as one participant did not fulfill the required three epochs after applying autoreject, the sample size of Dortmund Vital used in this study was *N* = 607. Participants met exclusion criteria including (1) neurological diseases, (2) cardiovascular, oncological, and eye diseases (3) psychiatric and affective disorders, (4) head injuries, head surgery, and head implants, (5) use of psychotropic drugs and neuroleptics, and (6) limited physical fitness and mobility. Furthermore, their study population was representative in terms of age distribution, genetics, cognitive performance parameters, and occupation, though it differed from the general German population in gender distribution and educational qualifications. EEG recordings were performed using a 64-channel elastic cap (10-20 system), a BrainVision Brainamp DC amplifier, and BrainVision Recorder software (BrainProducts GmbH). For our study, we used EEG data recorded prior to the two-hour block of cognitive experimental tasks.

##### AI-Mind

We used data from the AI-development set, which is a subset of the AI-Mind dataset [2]. This datasets contains participants with mild cognitive impairment, collected at Oslo University Hospital (Oslo, Norway), Helsinki University Hospital (Helsinki, Finland), Universidad Complutense de Madrid (Madrid, Spain), Catholic University of the Sacred Heart (Rome, Italy), and IRCCS San Raffaele (Rome, Italy). The EEG data used for this study was restricted to the first visit of the AI-Mind study. Furthermore, the p-tau 217 levels were also collected at the first visit, at the MCI stage. After excluding participants for whom a p-tau 217 measurement was unavailable, and restricting the dataset to only include participants with at least one available EEG recording from the eyes open condition and at least one available recording from the eyes closed condition, the sample sizes were Norway (*N*_*NO*_ = 152), Finland (*N*_*FI*_ = 66), Spain (*N*_*SP*_ = 158), and Italy (*N*_*IT*_ = 110), giving a total of *N* = 486 participants. It is important to note that this dataset does not represent the full AI-Mind study data, as a test hold-out set remains unavailable to AI research and development for a final unbiased evaluation of the AI-Mind tools. The EEG was recorded using 126 scalp electrodes prewired in an elastic cap (ANT neuro WaveguardTM), with electrode positioning following the 10-5 system derived from the extended international 10-20 system. Ethical approval for the AI-Mind study was granted to each data controller according to national guidelines. Please see [2] for details.

#### 2.2.3. Pre-processing

##### Input data

Our EEG pre-processing pipeline is shown in Figure 2. All pre-processing steps were implemented using tools from MNE-Python [47] and autoreject [49]. We employed HPO of certain steps of the pre-processing pipeline, resulting in multiple pre-processed versions of the input data. The first step of the pipeline was to remove the first 30 seconds of the data, as artefacts are more common in the start of a recording session. The data was band-pass filtered between (1 − 45)Hz, and after visual inspection of the power spectra, a notch filter at 50Hz was applied to the Dortmund Vital and AI-Mind datasets. Subsequently, the EEG data was epoched into four different copies, with duration *d* ∈ {5*s*, 10*s*, 20*s*, 30*s*} . For each copy, autoreject [49] was applied for channel repair and epoch removal. We used three epochs per participant, and participants which did not satisfy this criteria after autoreject was applied, was removed from the dataset. Interpolation/channel removal to the Standard 10-20 with 19, 32, 64, and 126 channels was employed ^4^. Furthermore, we used both spherical spline and biophysical interpolation. The EEG data was re-sampled to 90Hz, 135Hz, and 180Hz. Finally, average referencing was applied.

**Figure 2.**
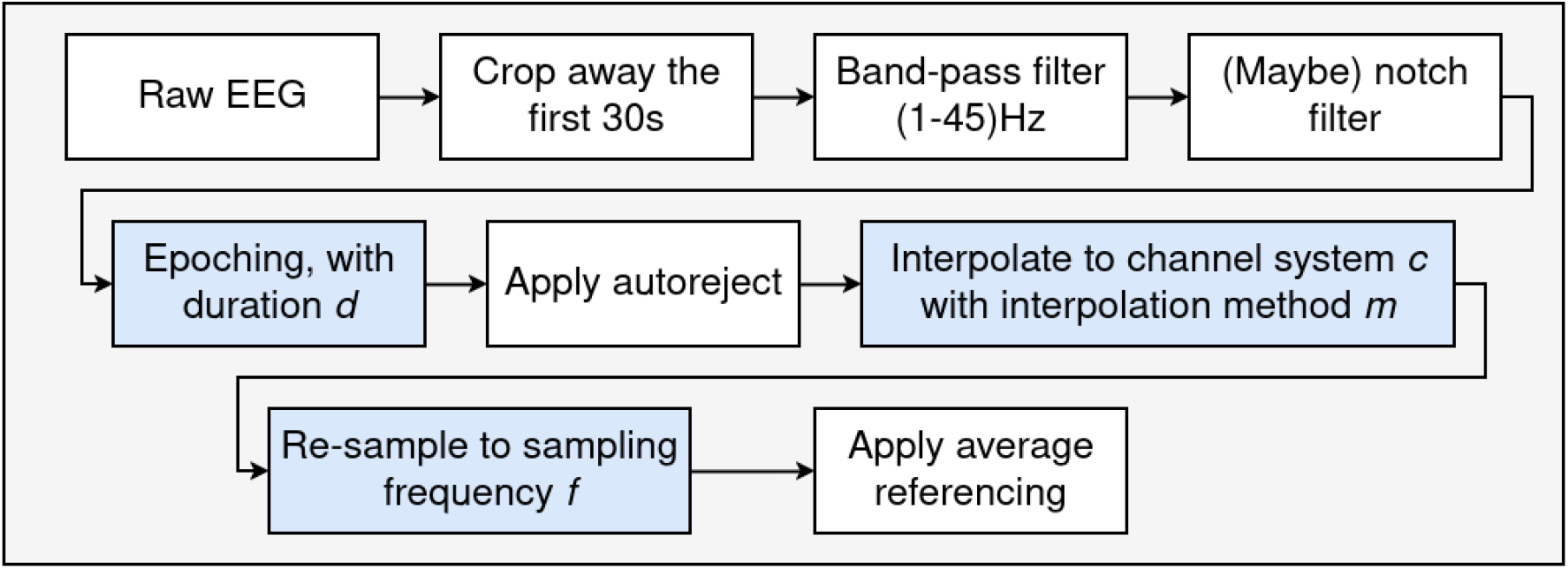
Pre-processing pipeline for creating multiple versions of the EEG data to be used as input to the DL models. The rectangles in blue indicates that the pre-processing step contains hyperparameters which are part of the hyperparameter optimisation. For these hyperparameters, we used the configuration space composed of input durations *d* ∈ {5*s*, 10*s*, 20*s*, 30*s* }, channel systems of the extended standard 10-20 system with number of channels in *c* ∈ {19, 32, 64, 126 }, *m* spline, biophysical, and *f* ∈ {90Hz, 135Hz, 180Hz }. Note, however, that neither channel system nor interpolation method were relevant if region-based pooling was used.

##### Pseudo-targets

The first steps of the pipeline for computing band power as pseudo-targets included data pre-processing. This was made by (1) cropping out the first 30 seconds and the last 15 seconds of data, (2) band-pass filtering between 1 and 45Hz, (3) applying a notch filter at 50Hz (4) re-sampling to 180Hz (5) epoching the data to 5-second non-overlapping segments (6) applying autoreject, and (7) applying average referencing. Subsequently, PSDs were computed per channel using the Welch method and averaged across epochs. Band-power was calculated by Simpson integration per channel and frequency band *ω* ∈ {*δ, θ, α, β, γ* }with *δ* = (1 − 4)Hz, *θ* = (4 − 7)Hz, *α* = (7 − 12)Hz, *β* = (12 − 30)Hz, and *γ* = (30 − 45)Hz . To avoid a high potential impact of bad channels, we used the median band power across channels. The bandpower values were multiplied by a factor of 10^12^, and transformed by a log-10 transformation. Finally, the pseudo-targets were z-normalised to obtain zero mean and unit standard deviation on the pretext training data.

##### Downstream targets

For the semi-supervised MTL-based optimisation (as well as baseline DL models, see Sec. 2.4), the downstream targets were z-normalised to obtain zero mean and unit standard deviation on the downstream training data. An inverse scaling was applied to the model outputs prior to computing metrics.

#### 2.3.4. Data splits and model selection

##### Supervised HPO

For training on the pretext tasks, we performed a single random split, reserving 20% of the subjects for validation. The epoch with the highest performance (as measured by *R*^2^ on the validation set) was selected, and its corresponding model state was used for the HPC evaluation. To evaluate a single HPC, we first partitioned the subjects from the downstream task into two sets, (*S*_*HPCE*_, *S*_*test*_), with 70% allocated to *S*_*HPCE*_ and 30% to *S*_*test*_. This partitioning remained fixed for all trials. To provide feedback to the HPO algorithm, we performed 50 random 80/20 splits of *S*_*HP CE*_. For each split, a linear regression model was trained using ordinary least squares (OLS) on 80% of the data and evaluated on the remaining 20%. The median *R*^2^ validation score across these splits was used as the signal to the HPO algorithm. *S*_*test*_ remained separated from the HPO process, ensuring an unbiased final evaluation of our methods.

##### Semi-supervised MTL

To obtain unbiased and comparable results with the supervised HPO strategy, we isolated the same test set as for supervised HPO. For each trial, model training and model state selection was performed by a single random split, reserving 20% of the subjects for validation. All epochs that were Pareto optimal (as measured by *R*^2^ on the validation set for both tasks) were stored, and the model state which had the highest smallest *R*^2^ score on the two tasks was used for the HPC evaluation.

#### 2.3.5. HP distributions

Figure 3 shows a shallow overview of the HPs and the corresponding HPDs used in our experiments. See Appendix A.1 for a complete overview. One guideline for selecting HPDs was to use fairly broad configuration spaces, which was motivated by the effective HP dimensionality often being considerably smaller than the number of HPs, and random search being robust in such scenarios [50]. A second guiding principle was to avoid configurations which vastly increased the runtime. While the majority of HPDs were pre-determined, some HPDs were adjusted due to high runtime and memory considerations. For this purpose, different HPDs were explored through trial and error on (1) age prediction using LEMON as the pretext dataset and Dortmund Vital for the downstream task, and (2) a fake downstream target which only contained noise, using LEMON and Dortmund Vital as pretext datasets and AI-Mind as the downstream dataset. All HPO was made automatic, with no opportunity for the researchers to make post-hoc adjustments to further refine the HPDs and initiate new experiments.

**Figure 3.**
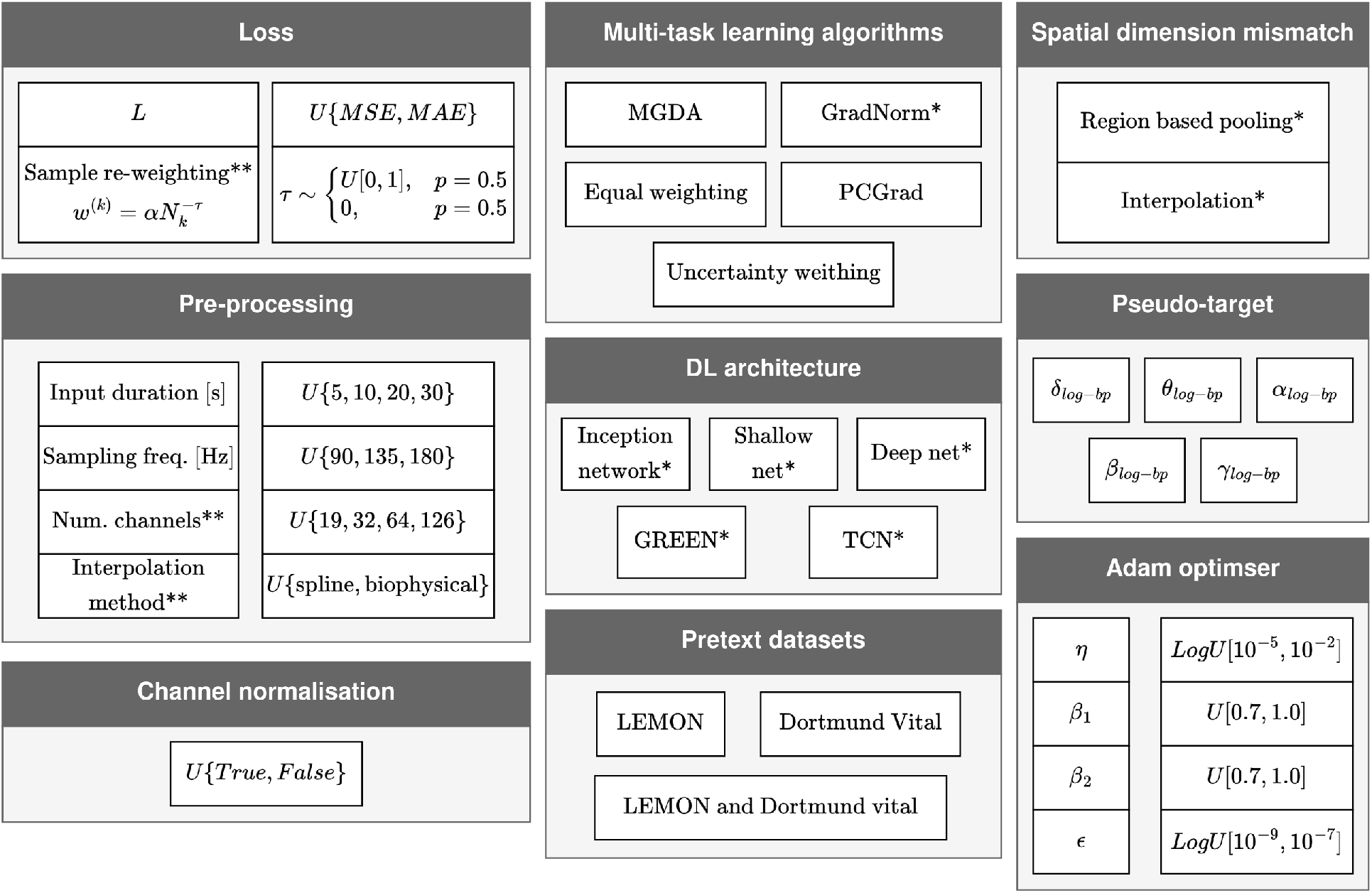
A shallow overview of the HPs and corresponding distributions. HPs marked with * contain additional HPs which are not shown in this figure. HPs marked with ** are conditional HPs. A complete overview is documented in Appendix A.1.

### 2.4. Experimental setups and baselines

This work was performed on the TSD (Tjenester for Sensitive Data) facilities, owned by the University of Oslo, operated and developed by the TSD service group at the University of Oslo IT-Department (UiO IT). We used the High Performance Computing resources of Colossus, the high performance computing cluster provided as part of TSD by UiO IT.

HPO was carried out using Optuna [51], setting both the ‘group’ and ‘multivariate’ arguments to True. All DL models were trained for 500 epochs, employing early stopping with a patience set to 50. For all models not employing semi-supervised MTL, patience refers to the number of consecutive epochs without improvement in the validation set’s *R*^2^ score before early termination is triggered. For models employing semi-supervised MTL, patience refers to the number of consecutive epochs without finding a new Pareto-optimal solution, as evaluated by the *R*^2^ score on the pretext validation set and the *R*^2^ score on the downstream validation set.

#### 2.4.1. Linear regression with band-power features

The first baseline was to employ a linear regression model to predict the target variables from band-power. The features were the same as the pseudo-targets, log band-power of {*δ, θ, α, β, γ* }frequency bands from the eyes open condition.

#### 2.4.2. Downstream training with DL

We used DL models to predict the downstream target without any form of pre-training, using the same HPDs as described in Sec. 2.3.5. Furthermore, we added ocular state as an HP, such that the input data could either be from eyes open or eyes closed. 500 random startup trials and 500 trials employing TPE was employed.

#### 2.4.3. Pretraining and fine-tuning

Although the primary aim of this study involved using the deviations from healthy expectation values as features for predictive modelling, we also investigated fine-tuning the models on the downstream task. The same HPDs as detailed in Sec. 2.3.5 were used. The loss function and the HPs of the Adam optimiser were considered separate for the pretext and downstream task, that is, their values were sampled independently. We used 500 random startup trials and 500 trials employing TPE.

#### 2.4.4. Supervised HPO: Simple linear regression with residuals

Here, we used the derived framework to learn expectation values on healthy EEG data, and using the residuals for predictive modelling on the downstream task. Only a single residual was used, and the frequency band that was used in generating the pseudo-target was considered an HP. We re-used the 500 random trials from Sec. 2.4.3 and continued optimising the HPs with 500 TPE-based trials.

#### 2.4.5. Supervised HPO: Multivariable linear regression with residuals

As an attempt to combine the learned residuals from Sec. 2.4.4 into a multivariable linear model, we combined the residuals that resulted from training with the different log band-power features as pseudo-targets. Here, one residual per frequency band was used for all trials. We added trials by (1) combining the highest performing HPCs (as evaluated by *R*^2^ on the downstream task) per frequency band, (2) 2500 trials following a sampling strategy which selects one random experiment per frequency band, and (3) 2500 trials following a sampling strategy which performs weighted sampling per frequency band. In (3), the weights were computed as softmax probabilities 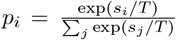, where *p*_*i*_ is the probability that the model of experiment *i* is selected, *s*_*i*_ is the validation *R*^2^ score on the downstream task, and *T* ∈ ℝ is a temperature scaling which is selected such that the top 25% is selected 75% of the times.

#### 2.4.6. MTL for learning residuals

We used HPO by 500 random trials followed by 500 trials employing multi-objective TPE [37, 51, 52].

### 3. Results

#### 3.1. Downstream predictive performance

Figure 4 shows the performance scores for all trials for the six different experiment-types from Sec. 2.4, including *R*^2^ and Pearson’s r. See Fig. A.6 for explained variance regression score, Spearman’s rho, and concordance correlation coefficient. After model selection, the results show negative *R*^2^ scores and weak correlations. Specifically, the performances scores were: linear regression with band-power features (*R*^2^ = − 0.02, Pearson’s r =0.18), downstream DL training (*R*^2^ = − 0.25, Pearson’s r = 0.14), pretraining and fine-tuning (*R*^2^ = − 0.29, Pearson’s r = 0.05), simple supervised HPO (*R*^2^ = − 0.10, Pearson’s r = 0.07), multivariable supervised HPO (*R*^2^ = − 0.05, Pearson’s r = 0.23), and semi-supervised MTL (*R*^2^ = − 0.24, Pearson’s r = 0.17). Additionally, even the highest performing models on the test set (which are clearly biased performance estimates) did not reach strong performance on the downstream task, suggesting that variance during model selection is not the limiting factor. In sum, neither our framework nor the baselines were able to reach high predictive performance for p-tau217 levels.

**Figure 4.**
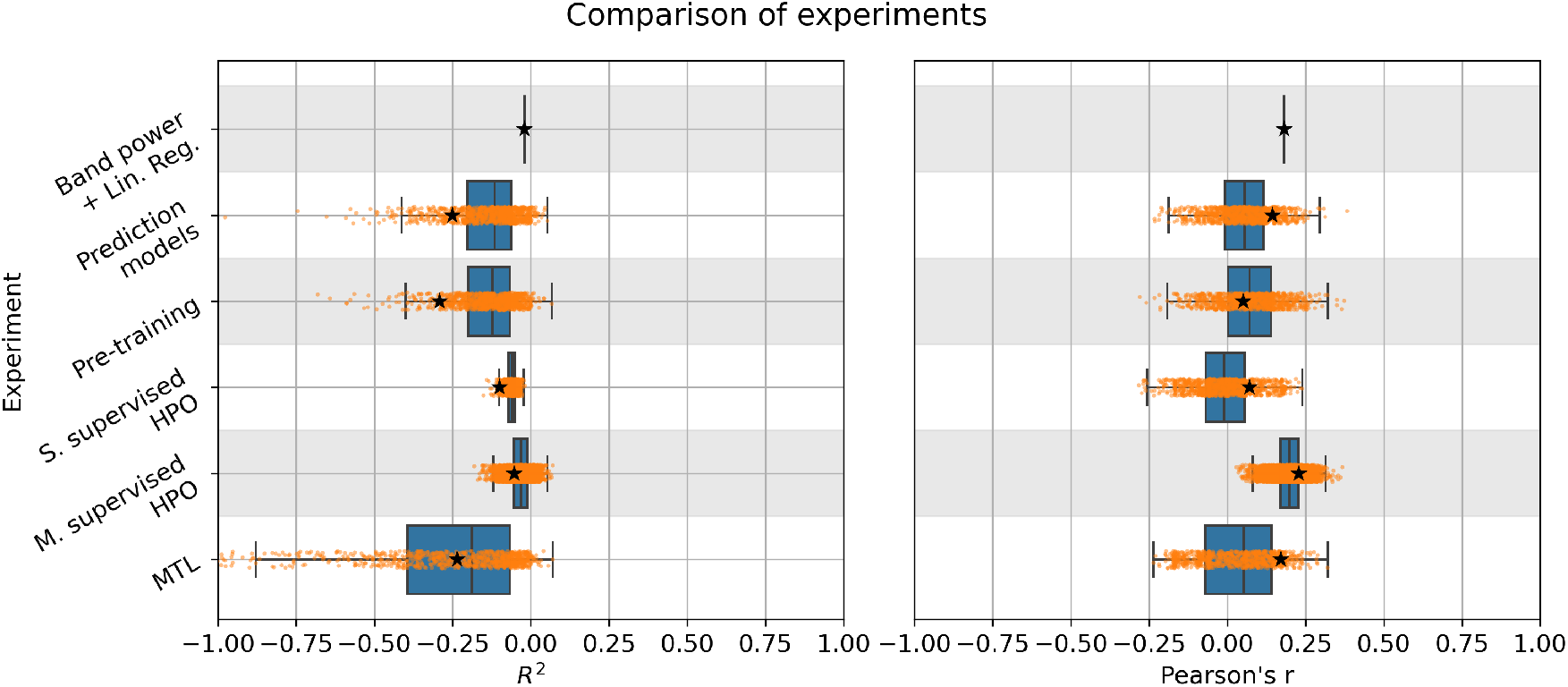
Test scores for all experiments and trials. The star indicates the test scores after model selection. Notably, all selected models obtained poor results, as indicated by negative *R*^2^ scores and weak correlations. See Fig. A.6 for similar plots with explained variance ratio score, Spearman’s rho, and concordance correlation coefficient

#### 3.2. Estimated Pareto front

Figure 5 shows the performance on the downstream task plotted against performance on the pretext task, for multiple regression metrics. The majority of solutions that were Pareto-optimal across all epochs and trials were obtained using band power from the *β* band. However, as the downstream performance scores were poor, this appears to result from high predictive values on the pretext task.

**Figure 5.**
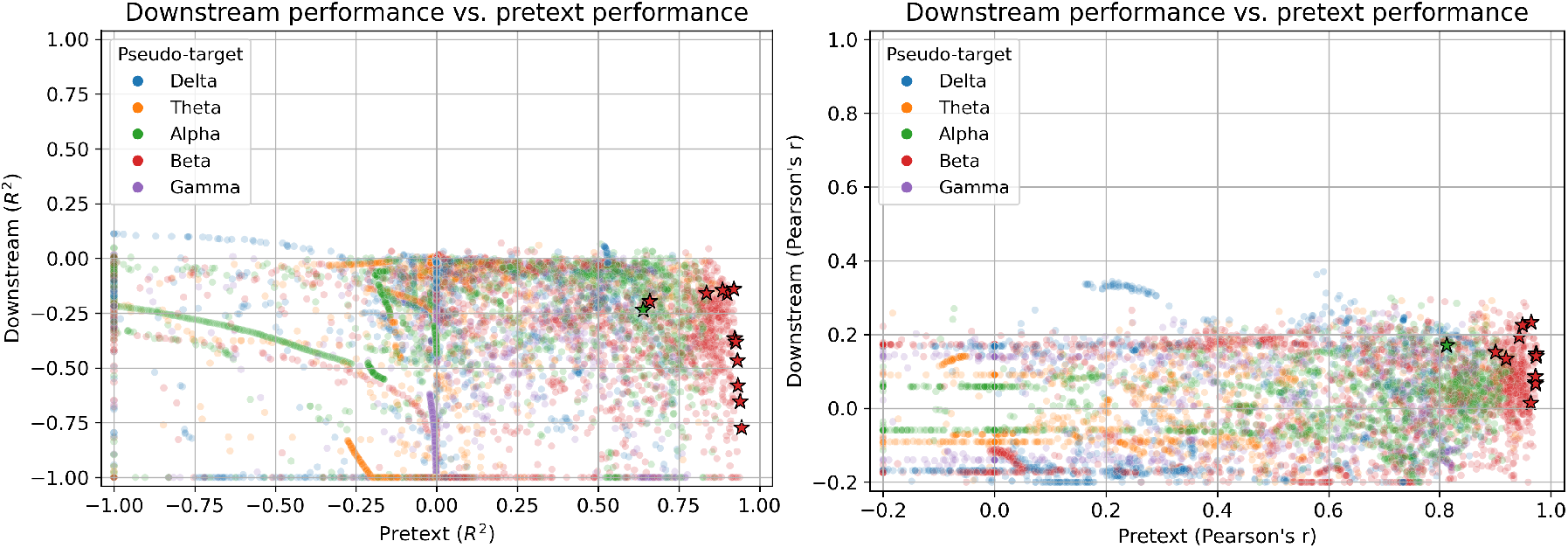
Performance scores on the downstream test set and pretext validation set for different metrics. Here, all Pareto optimal scores (as computed by *R*^2^ score on the downstream and pretext validation sets) per trial is plotted. The stars indicate that the solution was Pareto optimal as estimated on the validation sets across all trials, hence representing the estimated Pareto front during model selection. The downstream performance scores of the estimated Pareto front were poor, as indicated by low *R*^2^ scores and weak correlations. In contrast, the performance scores on the pretext task was substantially better, achieving high *R*^2^ scores and correlations. The majority of the estimated Pareto optimal solutions used the *β* power as pseudo-target. Performance scores below selected thresholds were truncated (*R*^2^ *<* − 1.0 and Pearson’s r *<* − 0.2). See Fig. A.7 for similar plots with explained variance ratio score, Spearman’s rho, and concordance correlation coefficient.

The performance scores on the pretext task were high, ranging from 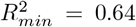 to 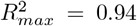 and Pearson’s r_*min*_ = 0.81 to Pearson’s r_*max*_ = 0.97 for the estimated Pareto-optimal solutions. On the downstream task, these models obtained performances scores ranging from 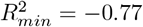 to 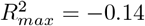 and from Pearson’s r_*min*_ = 0.01 to Pearson’s r_*max*_ = 0.23. Therefore, high performance scores on the pretext task did not translate into high downstream performance scores. This suggests that the model was able to capture healthy patterns of electrophysiological reactivity, but that deviations from these healthy patterns were not predictive of neurodegeneration.

## 4. Discussion

This work designed and tested a novel SSL-based framework which learns healthy ocular state transitioning patterns. Furthermore, it used two optimisation strategies which explicitly aim to find the healthy transitioning trends which also breaks for pathological data, to promote residuals of high predictive value in pathology decoding. The framework is conceptually simple - computing deviations from learned expectation values. This may increase its impact for biomarker discovery, and be a step towards more neurophysiologically grounded DL modelling. A connection to the pathology is indeed of high importance for developing biomarkers [53, 54].

We hypothesised that the main advantage of our framework over traditional DL-based predictive modelling was its potential biological insights, but unlikely to reach traditional DL performance scores. From a technical perspective, imposing our framework represents a major constraint for empirical risk minimisation, hence, the baselines which applied DL models directly on the downstream task were unlikely to be surpassed, but in practice serve as an upper bound instead. Additionally, we hypothesised that an advantage to traditionally computed reactivity was that the state transitions could be learned in a data-driven manner, potentially discovering reactivity patterns that are currently unknown and recognised. Indeed, alpha reactivity stands out as the most documented reactivity pattern for dementia-related cognitive health [14, 13, 55]. However, to maintain high exploration in our experiments, we did not explicitly filter out the alpha signal, and we also considered other frequency band-powers as pseudo-targets. Instead, we allowed the DL models to perform such feature extraction through data-driven learning. Additionally, results from computer vision has shown that the first layer of deep neural networks occasionally learns features similar to Gabor filters [56], and such filters are also explicitly modelled by the GREEN architecture [57].

Our work is distinct from traditional SSL methods because our framework aims to learn patterns of healthy transitioning patterns and use the deviation from expectation value as a biomarker, rather than learning latent vector representations. Common pretext tasks for SSL applied to data EEG data, however, typically focus on learning representations through contrastive or predictive objectives, such as masking, shuffling, time- and frequency contrasting, and signal transformation recognition [31, 30, 26, 27, 28, 29, 24, 21, 22, 23, 25]. While effective for downstream classification [35], these approaches do not explicitly model normative dynamics nor capture deviations that may signal pathology. In contrast, our method leverages the actual-to-predicted residual as a measure of abnormality, offering interpretable markers that are better tied to physiological state changes.

Adapting DL techniques for EEG biomarker discovery is an interesting area of research. A notable example is the GREEN architecture, which incorporates learnable Gabor wavelets and Riemannian geometry to embed inductive biases directly into the network design [57]. A key distinction between GREEN and our approach lies in how and where inductive bias is introduced: GREEN modifies the model architecture itself, whereas our framework embeds neurophysiological priors through reactivity-inspired pretext tasks during the self-supervised training phase and enforces residual-based biomarkers. Importantly, as demonstrated in this study, architectural innovations and novel training-phase strategies are not mutually exclusive and may be combined to enhance both biomarker interpretability and predictive performance.

Despite thorough HPO, our developed framework did not achieve high predictive performance for p-tau217 levels. However, by using the same HPDs (and including ocular state as an HP) and employing downstream training only, these baselines obtained similar poor results. This suggests that the lack of predictive performance may not be due to the framework itself, but from limitations in the data. Specifically, resting-state EEG may not encode signal components that are reliably associated with the pathological processes approximated by p-tau217 levels at the MCI stage. Furthermore, the use of p-tau217 as a surrogate marker introduces additional complexity, as it is not a direct neural signal but rather a downstream biomolecular correlate of neurodegeneration.

### 4.1. Connections to reactivity

Reduced reactivity in posterior alpha rhythms during the transition from eyes-closed to eyes-open conditions has been linked to cognitive pathology. For instance, reduced posterior alpha reactivity has been associated with Parkinson’s disease dementia [14], and similar reductions have been observed in patients with dementia due to Lewy body and Alzheimer’s disease, compared to healthy controls [13, 55]. Notably, alpha reactivity has also demonstrated predictive value for amyloid-beta positivity beyond established clinical measures [58], and some evidence suggests that EEG reactivity is preserved in healthy ageing [59]. In addition, alpha reactivity appears to be sensitive to vestibular dysfunction [60], and reduced alpha reactivity has been linked to compromised integrity of the cholinergic system [61, 62].

These findings highlight the relevance of alpha reactivity as a biomarker, but also reveal a reliance on hand-crafted EEG features. In contrast, our framework offers a data-driven approach for learning transition patterns between brain states, such as the eyes-closed to eyes-open condition. By using a self-supervised objective that models normative reactivity and captures deviations as residuals, our method offers a flexible way to discover latent, potentially unknown patterns that may correspond to underlying pathology. This is further enhanced by optimisation strategies that explicitly encourage the discovery of transition patterns that are both representative of healthy neural functioning and sensitive to pathological deviations. This approach complements existing physiologically informed biomarkers and opens a path toward improved data-driven biomarker discovery.

### 4.2. MTL for learning relevant residuals

MTL is traditionally motivated by goals such as improving predictive performance, enhancing generalisation, learning from partially annotated data, and reducing computational cost [63, 64, 65, 66, 67, 68, 69]. The core idea is that jointly solving multiple tasks using a shared encoder can lead to better representations than training separate models for each task.

While our objective differs from these typical motivations, algorithms from the MTL literature are readily applicable. Specifically, we employed MTL not to jointly solve multiple downstream tasks, but to guide the model toward producing interpretable biomarkers that are also predictive of pathological outcomes. Rather than enhancing task performance directly, the goal is to shape residuals in a way that captures clinically relevant deviations from normative brain dynamics.

The idea behind formulating our problem using both a pretext and a downstream objective (and applying appropriate gradient-based algorithms from MTL literature) may intuitively be understood through a thought experiment, using alpha power as an example pseudo-target. Imagine that there are multiple features derived from eyes-closed recordings which may be used to predict the alpha power during eyes open, including, e.g., alpha power, theta/delta power ratio, beta/gamma cross-frequency correlations. Among all these possible transitioning patterns, one is not necessarily interested in the feature(s) which obtains the smallest prediction error on this pretext task, but rather to look for the transitioning patterns which are sensitive to abnormalities due to the presence of pathology. Using machine learning with hand-crafted features, this may be approached by domain knowledge and hypothesis-driven research, where the features are specified by the researcher. However, as one has limited control over which features are learned by DL models, imposing such feature-restrictive inductive biases is often not feasible. We therefore argue that for DL models, it is particularly crucial that the optimisation strategies are aligned with our actual goal. The two-task formulation makes the problem fundamentally more challenging, as we now have two optimisation problems without any clear way of comparing different Pareto optimal solutions. However, we argue that solely solving the pretext task is an incomplete optimisation strategy^5^, and that the added complexity represents a more realistic reflection of biomarker discovery, where both biological groundedness and predictive performance must be balanced.

Importantly, this approach is not limited to the reactivity- and SSL-based framework developed in this study. The underlying principle, using MTL to increase the predictive value of residuals, can be extended to any residual-based biomarker. Its most straightforward application is to formulate age prediction as a multi-objective optimisation problem, in which the model simultaneously minimises age prediction error and maximises the association between the age gap and a target pathology. Following this formulation (as mathematically expressed in Eqs. 6 and 7), MTL algorithms may help uncover age-related trends that are particularly sensitive to disease processes. This broader perspective suggests that MTL can be a powerful tool for enhancing residual-based biomarkers across diverse domains of EEG and biomedical analysis.

### 4.3. Future work

Our framework may be generalised beyond ocular reactivity. As reactivity has been explored for different purposes than dementia research, it would be interesting to test our methods on such data. An interesting domain of application may be consciousness impairment, where preserved reactivity has been linked to an increased odds of survival [15]. In consciousness impairment, other stimuli such as auditory and nociceptive stimuli has been used [15], which impose different requirements for the datasets used in the pretext task. Additionally, reactivity has shown high predictive value of neurological outcome after cardiac arrest [16], and would therefore be of high interest for further investigation of our approach. We highlight that, although our framework was formulated in the cross-ocular state setting, the extension to other EEG recording paradigms is trivial. By re-defining *O*_1_ as the ‘relaxed state’ and *O*_2_ as the ‘stimulated state’, such as *O*_2_ being the state in which auditory stimuli was applied, our methods are still valid and applicable.

Although our experiments only implemented DL-based models for the SSL-based framework, this is not a requirement for using our proposed methods. ML techniques based on hand-crafted feature extraction are indeed compatible with our framework, and may in particular be beneficial for explainability, further enhancing the derived neurophysiological insight. Alternatively, inspecting explainable DL models such as GREEN [57] could be of high interest.

In both our framework and our experiments, the pseudo-targets were pre-selected. It is conceivable, however, that the pseudo-target may benefit from being learned as part of the gradient-based optimisation. That is, to use 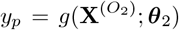 and add ***θ***_2_ to the optimisation in Eqs. 6 and 7. For maintaining the neurophysiological insight however, this approach requires that the model *g*(·) is sufficiently interpretable, so that an interpretation of the expectation value and corresponding residual can be made.

## 5. Conclusion

In this study, we designed a novel self-supervised learning based framework for modelling patterns of healthy transitions from eyes-closed to eyes-open resting-state EEG. The residual, representing deviation from healthy expectation value, is used as a candidate biomarker. We hypothesised that this framework could yield more biologically interpretable outputs compared to traditional DL methods, and that currently unknown reactivity patterns could be learned in a data-driven manner. To promote residuals that are sensitive to pathological abnormalities, we applied two optimisation strategies which involves maximising the predictive value of the residual. Neither our proposed framework nor our baselines achieved satisfactory predictive performance for ptau-217 levels, which were used as a surrogate marker of cognitive health. Therefore, the main limitation is likely due to insufficient signal components in the EEG that are strongly associated with the pathological processes approximated by p-tau217 levels at the MCI stage. Hence, this study represents a methodological contribution to the development of interpretable, transition-based EEG biomarkers through self-supervised learning, but its clinical utility remains empirically unverified. Future studies with more sensitive downstream markers are needed to validate the utility of our framework in clinical decision-making.

## Acknowledgments

This project has received funding from the European Union’s Horizon 2020 research and innovation programme under grant agreement No 964220. This publication reflects views of the authors and the European Commission is not responsible for any use that may be made of the information it contains.

## Author contributions

**Thomas Tveitstøl**: Conceptualization, Data curation, Formal analysis, Investigation, Methodology, Software, Visualization, Writing – original draft, Writing – review & editing **Mats Tveter**: Investigation, Writing – review & editing **Christoffer Hatlestad-Hall**: Writing – review & editing **Hugo L Hammer**: Supervision, Writing – review & editing **Ira R J Hebold Haraldsen**: Funding Acquisition, Writing – review & editing

## Appendix

### Appendix A.1. Hyperparameter distributions

In this section, we document all the HPDs that were used. Note that the sampling distributions only hold during the initial random search, not for the trials employing TPE.

#### Appendix A.1.1. Datasets for the pretext task

The datasets that were used for training on the pretext tasks were either LEMON, Dotmund Vital, or both, sampled with equal probabilities.

#### Appendix A.1.2. Pre-processing

The HPs involved in the pre-processing of the data were sampling rate and input length. Additionally, the selection of channels system and the interpolation method were conditional HPs constrained to trials employing interpolation for handling the varied electrode configurations. The corresponding HPDs are detailed in table A.1.

**Table A1.** Sampling distributions for the pre-processing steps. *U*{*a*_1_, *a*_2_, …, *a*_*n*_} denotes uniform sampling over the discrete set {*a*_1_, *a*_2_, …, *a*_*n*_}

| HP | Distribution |
| --- | --- |
| Sampling rate | $U\{90\text{Hz}, 135\text{Hz}, 180\text{Hz}\}$ |
| Input length | $U\{5s, 10s, 20s, 30s\}$ |
| Channel system* | $U\{\text{Standard19}, \text{Standard32}, \text{Standard64}, \text{Standard126}\}$ |
| Interpolation method* | $U\{\text{spline}, \text{MNE}\}$ |

\* These HPs were only relevant when interpolation was used for handling the varied electrode configurations.

#### Appendix A.1.3. DL architectures

We considered five DL architectures, which were Inception network [70], ShallowFBCSPNet [71], Deep4Net [71], TCN [72], and GREEN [57]. Below, we provide a brief overview of each architecture. For complete descriptions, see the original works. In addition, we outline the sampling distributions of their HPs as used in our experiments.

For all architectures, the channels were normalised to 0 mean and unit standard deviation before its forward pass, with 50% probability.

##### Inception network [70]

The main building block of Inception network is the Inception module, which performs convolution with kernels of different sizes and aggregates the resulting feature maps by concatenation. Three Inception modules in series builds a single residual block. Inception network was originally composed of two such residual blocks, connected by a shortcut layer.

The number of residual blocks was sampled from the logarithmic distribution *d* ∼ *LogU* _ℕ_ [1, 12]. Furthermore, the number of convolutional filters of the inception modules was sampled from *f* ∼ *LogU* _ℕ_ {4, 64}.

##### Deep4Net [71]

Deep4Net consists of four convolutional-max pooling blocks. The first block includes both temporal and spatial filters, while the subsequent three blocks use convolution and max-pooling, with the number of convolutional units doubling for each successive block.

Padding on the last three convolutional layers was added. This was added to allow for a smaller number of input time steps. The kernel length for all convolutions, with the exception of the spatial filter in the first convolutional block, was fixed equal and sampled from *l* ∼ *U* _ℕ_ [5, 15]. The number of filters for the temporal and the spatial convolutional layers in the first convolutional block was fixed equal and sampled from *f*_1_ ∼ *U* _ℕ_ [10, 40]. The subsequent layers were computed as *f*_*i*_ = 2*f*_*i*−1_∀*i* ∈ {2, 3, 4}. Drop-out on the last layer was applied, and sampled from drop-out ∼ *U* [0, 0.5].

##### ShallowFBCSPNet [71]

ShallowFBCSPNet was inspired by the filterbank common spatial pattern pipeline. Similar to Deep4Net, its first two layers comprise separate temporal and spatial convolutional layers. These are followed by a squaring non-linearity activation function, a mean pooling layer, and a logarithmic activation function.

The kernel length of the temporal convolution in the first layer was sampled from *l* ∼ *U* _ℕ_ [5, 45]. As for Deep4Net, the number of filters for the temporal and the spatial convolutional layers in the first convolutional block was fixed equal. The number of filters was sampled from *f*_1_ ∼ *U* _ℕ_ [20, 60], the number of strides of the mean pooling layer was *t* ∼ *U* _ℕ_ [5, 25], and the kernel length of the pooling layer was computed as 5*t*. Drop-out on the last layer was sampled from drop-out ∼ *U* [0, 0.5].

##### TCN [72]

TCN employs dilated convolutional 1D filters to obtain exponentially large receptive fields. Furthermore, it uses residual modules which consist of, repeated twice, (1) a dilated causal convolution with weight normalisation, (2) Relu activation function, (3) and a spatial drop-out layer operating per channel. To ensure that the input and output tensors can be added together at the skip connection layer, a convolution with a kernel size of 1 × 1 is applied, if needed, to the input tensor to match the feature dimension of the output tensor.

Originally, the TCN architecture makes one prediction per time step. To obtain a scalar output, we added a global average pooling layer in the temporal dimension prior to the output layer. The dilation factor was fixed at *d* = 2. The number of residual blocks was sampled from *U* _ℕ_ [1, 5]. The number of filters per residual block was sampled from *LogU* _ℕ_ [4, 128]. Furthermore, the kernel size in the temporal dimension was sampled from *U* _ℕ_ [3, 8]. Finally, the drop-out rate was sampled from *U* [0, 0.5].

##### GREEN [57]

The Gabor Riemann EEGNet (GREEN) is a recently developed architecture which integrates domain knowledge by combining DL with spectral analysis and geometric operations. The first module of the network is *parametrised convolution*, which are wavelets on the form 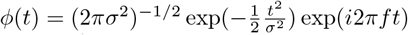, where *σ* ∈ ℝ^+^ is the standard deviation of the Gaussian window and *f* ∈ ℝ^+^ is the frequency of the complex sinusoid. Here, *σ* and *f* are treated as trainable parameters. The second module of GREEN is the *pooling layer*, which transforms the wavelet-transformed signals to non-temporal features. The original work implemented pooling layers which compute (1) the real covariance matrix, (2) the real covariance matrix with cross-frequency interactions, (3) pairwise phase-locking values, (4) cross-frequency phase-locking value, (5) a combination of real covariance matrix and pairwise phase-locking, and (5) a combination of cross-frequency covariance matrix and cross-frequency phase-locking value. To mitigate potential numerical instabilities due to overestimation of the range of eigenvalues in the sample covariance matrices, a covariance shrinkage operation was applied as a subsequent step. Furthermore, a BiMap layer was implemented to mitigate issues with rank deficient covariance matrices. This is followed by a rectified eigenvalue (ReEig) layer, and a logarithm mapping. Finally, a fully connected module is applied.

The number of complex wavelets used in the parameterised convolution module was sampled from the distribution *LogU* _ℕ_ [1, 10]. For initialisation of *σ* and *f*, we used either the Morlet parametrisation or random initialisation, sampled with equal probabilities. The lower and upper bound of the initialisations of *f* where *f*_*min*_ = 1, *f*_*min*_ = 2^5.5^. The upper and lower bound of the full width half maximum (FWHM) where *FWHM*_*max*_ = 2^1^, *FWHM*_*min*_ = 2^−4.5^. The strides of the convolution was sampled from *U* _ℕ_ [1, 5]. The kernel width was sampled from *U* [2*s*, 4*s*].

For the pooling layer, we used either the real covariance matrix, cross-frequency covariance matrix, pairwise phase-locking value, cross-frequency phase-locking value, a combination of real covariance matrix and pairwise phase-locking, or a combination of cross-frequency covariance matrix and cross-frequency phaselocking value, sampled with equal probabilities. When using either two of the phase-locking pooling layers, we sampled the ‘reg’ HP from reg ∼ *LogU* [10^−7^, 10^−5^]

In the shrinkage layer, we initialised the shrinkage parameter by sampling from the distribution *U* [− 5, − 1]. For the BiMap layer, the output dimension was computed as [*x* ·(#input channels − 1)] where [·] is the rounding operator, with *x* ∼*U* [0.5, 1.0].

For the ReEig layer, the eigenvalue regularisation parameter was sampled from *LogU* [10^−5^, 10^−3^]. In the LogMap layer, we used either identity or the running log-Euclidean mean as reference, sampled with equal probabilities. The momentum for in the update rule (Eq. 7 in [57]) was sampled from *U* [0.8, 1.0]. If the matrix passed to the LogMap layer was ill-conditions, hence failing to compute eigenvalues, the trial was pruned.

The number of units for the input layer of the fully connected module was sampled from *U* _ℕ_ [16, 128], whereas the number of hidden layers was sampled from *U* _ℕ_ [1, 3]. The number of units for a given layer was computed as half of its prior. Finally, the drop-out rate was sampled from *U* [0, 0.5].

#### Appendix A.1.4. Varied electrode configurations

To allow for varied electrode configurations with different numbers of channels, we used interpolation or region-based pooling (RBP) [73], sampled with equal probabilities. When interpolation was used, the interpolation method and channel system was sampled from the distributions described in table A.1.

When using RBP, the algorithm for splitting the montage into different regions was the same as in the original work. This includes to repeatedly compute the centroid of 2D projections of channel positions, and generate *k* angles which partitions the current region such that the newly formed regions contains the same number of electrodes. To ensure that all channel systems were compatible with all montage splits, we used the intersection of electrodes to fit the montage splits. The split vectors, which determines all the *k*- values per region split, were **k**_1_ = (2, 2, 2, 2, 2, 2, 2)^T^, **k**_2_ = (3, 3, 3, 3, 3, 3, 3, 3)^T^, **k**_3_ = (2, 3, 2, 3, 2, 3, 2, 3, 2)^T^, **k**_4_ = (4, 3, 2, 3, 4, 3, 2, 3, 4)^T^. For a single montage split, the split vector was sampled with equal probabilities from the discrete set {**k**_1_, **k**_2_, **k**_3_, **k**_4_}. The stopping criterion, which prevents the generation of a new split if the number of electrodes is below this threshold, was sampled from *U* _ℕ_ [1, 6]. Finally, the total number of montage splits was sampled from *m* ∼ *LogU* _ℕ_ [1, 8].

We considered three pooling methods for aggregating channels within a region; (1) averaging, (2) channel attention, and (3) channel attention with a head region. As in the original work, the attention mechanism relied on feature extraction similar to ROCKET [74], which involved applying random convolutional kernels followed by computing the mean and maximum value of the feature maps. We allowed for the use of multiple *pooling modules*, which uses parameter sharing across the montage splits it operates on. The use of a single pooling module was employed with a 50% probability. Otherwise, the number of pooling modules was computed as *p* = *max*(1, [*xm*]), with *x* ∼ *U* [0, 0.5]. The number of montage splits per pooling module was allocated by initialising *p* bins with unit cardinality and incrementing a randomly selected bin *m* – *p* times. The size of the *i*-th bin was used as the number of montage splits for the *i*-th pooling module. When using pooling modules with a channel attention mechanism, the number of kernels was sampled from *LogU* _ℕ_ [10, 500], and their maximum receptive field was sampled from *LogU* _ℕ_ [15, 100]. For channel attention with a head region, the dimensionality of the search vector was sampled from *LogU* _ℕ_ [4, 32]. Finally, the parameter sharing of the embedding functions in Eqs. (3-4) and (6-7) in [73] were used with a 50% probability.

#### Appendix A.1.5. Loss and optimiser

The Adam optimiser was used for training the DL models on their pretext tasks. The learning rate was sampled from *η* ∼ *LogU* _ℕ_ [10^−5^, 10^−2^]. Furthermore, the beta values were sampled from *β*_1_ ∼ *U* [0.7, 1] and *β*_2_ ∼ *U* [0.7, 1], and the epsilon value for numerical stability was sampled from *ϵ* ∼ *LogU* _ℕ_ [10^−9^, 10^−7^]. The batch size was fixed to 64.

The loss function was *MAE* or *MSE*, sampled with equal probabilities. If the training included both LEMON and Dortmund Vital, a sample re-weighting was employed with a 50% probability. This included weighting the subjects according to the size of the dataset they belong to by 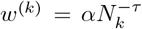, where *τ* is an HP which determines the power of re-weighting (*τ* = 0 corresponds to equal weighting of all subjects, while *τ* = 1 corresponds to equal weighting of all datasets), *N*_*k*_ ∈ ℕis the training sample size of the *k*-th dataset and *α* is a normalisation constant which ensures weighted averaging [75]. For the trials employing this re-weighting, *τ* was sampled from the distribution *τ* ∈ *U* [0, 1].

#### Appendix A.1.6. Algorithms for multi-task learning

In semi-supervised MTL, the two tasks were jointly optimised using (1) equal weighting of the loss functions, (2) GradNorm [38], (3) MGDA [39, 40], (4) PCGrad [41], or (5) uncertainty weighting [42], with equal probabilities.

In GradNorm, we used stochastic gradient decent on the weights *w*_*i*_ with a learning rate sampled from *LogU* [10^−3^, 10^−1^]. Furthermore, the *α* HP which is related to the training rate balance between tasks was sampled from *α* ∼ *U* [0, 3]

## Appendix A.2. Additional performance scores

Fig A.6 shows performance scores for all trials for the six different experiment-types from Sec. 2.4, including explained variance regression score, Spearman’s rho, and concordance correlation coefficient. After model selection, the performance scores were linear regression with band-power features (explained variance = 0.03, Spearman’s rho = 0.17, concordance correlation = 0.08), downstream DL training (explained variance = − 0.25, Spearman’s rho = 0.12, concordance correlation = 0.13), pretraining and fine-tuning (explained variance = − 0.20, Spearman’s rho = 0.02, concordance correlation = 0.04), simple supervised HPO (explained variance = − 0.04, Spearman’s rho = −0.01, concordance correlation = 0.04), multivariable supervised HPO (explained variance = 0.01, Spearman’s rho = 0.17, concordance correlation = 0.16), and semi-supervised MTL (explained variance = −0.09, Spearman’s rho = 0.14, concordance correlation = 0.12).

**Figure A6.**
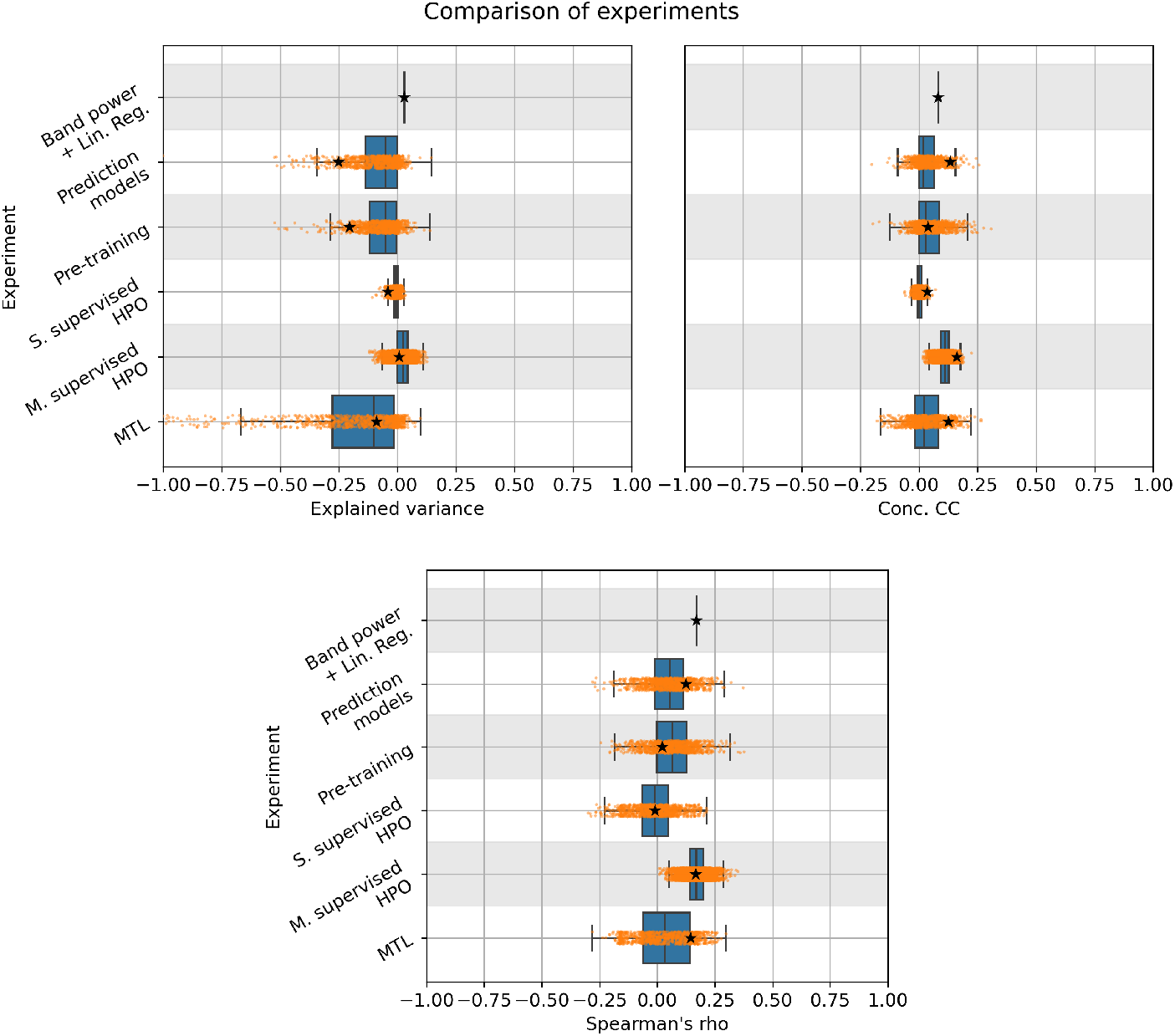
Test scores for all experiments and trials. The star indicates the test scores after model selection. Here, explained variance refers to the offset-invariant explained variance regression score as computed by the ‘explained_variance_score’ func- tion in sklearn [76]. Notably, all selected models obtained negative R^2^ scores, low explained variance regression scores, and weak correlations.

Fig. A.7 shows the performance on the downstream task plotted against performance on the pretext task, for explained variance regression score, Spearman’s rho, and concordance correlation coefficient. The Pareto-front, representing the scores of the models that were Pareto-optimal across all epochs and trials on validation *R*^2^ scores, obtained pretext test scores ranging from explained variance_*min*_ = 0.64 to explained variance_*max*_ = 0.94, Spearman’s rho_*min*_ = 0.72 to Spearman’s rho_*max*_ = 0.97, and concordance correlation_*min*_ = 0.79 to concordance correlation_*max*_ = 0.97. On the downstream task, the test performance scores ranged from explained variance_*min*_ = − 0.53 to explained variance_*max*_ = 0.04, Spearman’s rho_*min*_ = − 0.08 to Spearman’s rho_*max*_ = 0.25, and concordance correlation_*min*_ = 0.01 to concordance correlation_*max*_ = 0.18.

**Figure A7.**
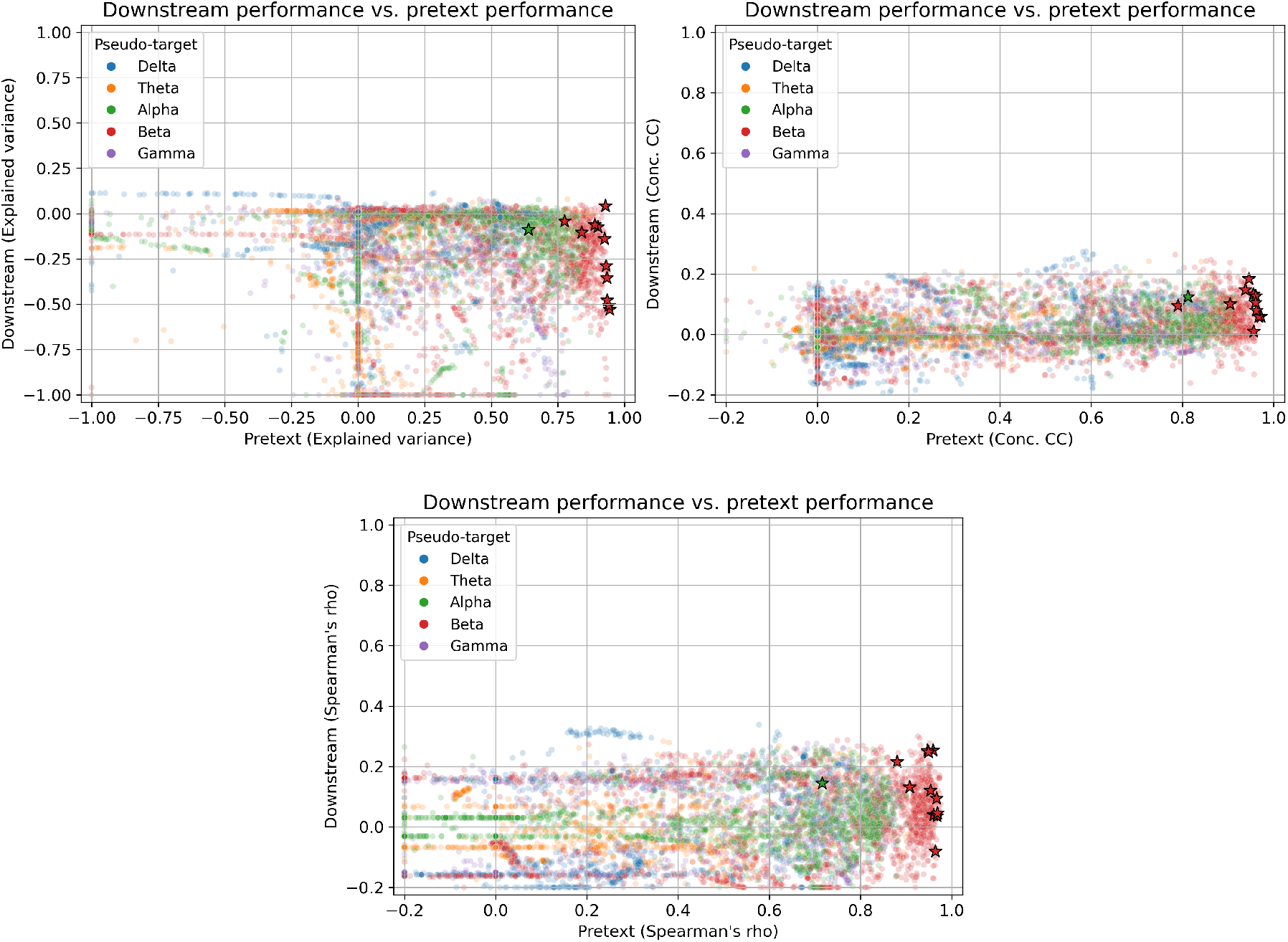
Performance scores on the downstream test set and pretext validation set for different metrics. Here, all Pareto optimal scores (as computed by R^2^ score on the downstream and pretext validation sets) per trial is plotted. The stars indicate that the solution was Pareto optimal as estimated on the validation sets across all trials, hence representing the estimated Pareto front during model selection. Performance scores below selected threshold were truncated (Explained variance < −1.0, concordance correlation < −0.2, and Spearman’s rho < −0.2).

## Footnotes

2 Our framework is not restricted to DL, but can be used with any prediction model. For consistency with our experiments, however, we will assume a DL model throughout the methods section.

3 Setting *O*_1_ = *O*_2_ removes the “cross-ocular” aspect but is unproblematic for those interested in learning within-ocular-state expectation values.

4 when region based pooling was used for handling varied electrode configurations, no interpolation was employed

5 in our framework, solving the downstream task without solving the pretext task would also be incomplete, as it would imply that the residual cannot meaningfully be interpreted as a deviation from an expectation value, leading to a lack of biological insight.

